# Development of highly multiplex targeted proteomics assays in biofluids using the Stellar mass spectrometer

**DOI:** 10.1101/2024.06.04.597431

**Authors:** Deanna L Plubell, Philip M. Remes, Christine C. Wu, Cristina C. Jacob, Gennifer E. Merrihew, Chris Hsu, Nick Shulman, Brendan X. MacLean, Lilian Heil, Kathleen Poston, Tom Montine, Michael J. MacCoss

## Abstract

The development of targeted assays that monitor biomedically relevant proteins is an important step in bridging discovery experiments to large scale clinical studies. Targeted assays are currently unable to scale to hundreds or thousands of targets. We demonstrate the generation of large-scale assays using a novel hybrid nominal mass instrument. The scale of these assays is achievable with the Stellar^TM^ mass spectrometer through the accommodation of shifting retention times by real-time alignment, while being sensitive and fast enough to handle many concurrent targets. Assays were constructed using precursor information from gas-phase fractionated (GPF) data-independent acquisition (DIA). We demonstrate the ability to schedule methods from an orbitrap and linear ion trap acquired GPF DIA library and compare the quantification of a matrix-matched calibration curve from orbitrap DIA and linear ion trap parallel reaction monitoring (PRM). Two applications of these proposed workflows are shown with a cerebrospinal fluid (CSF) neurodegenerative disease protein PRM assay and with a Mag-Net enriched plasma extracellular vesicle (EV) protein survey PRM assay.

## INTRODUCTION

An important lesson from the last decade is that immunoassays, or any other assay that relies on affinity reagents, are far from a perfect solution to the protein quantification problem. While the use of immunoassays has historically been viewed as the necessary endpoint for putting a protein assay into clinical practice, the negatives are beginning to outweigh the strengths. Multiplex affinity assays have been widely used in applications of quantitative protein measurements across 1000s of plasma or serum samples. While increased multiplexing to 100s or 1000s of proteins can minimize sampling bias and provide a more comprehensive sampling of the proteome, it can present significant challenges. Despite the promised depth of coverage of low abundance proteins in both SOMAScan and Olink**®**, LC-MS is still the gold standard method for providing selective protein measurements^1–3^. Mass spectrometry has always outperformed affinity-based methods when specificity is the priority^4–6^. In addition, mass spectrometry’s ability to measure multiple peptides per protein provides an opportunity to capture signals from different proteoforms^7^.

Because of limitations of affinity-based protein assays, targeted mass spectrometry assays were viewed as a promising alternative to immunoassays in the clinical lab. Early methods for performing targeted protein quantitation were performed using an acquisition strategy known as selected reaction monitoring (SRM) on a triple quadrupole mass spectrometer. This method was powerful as this hardware was common in mass spectrometry labs focused on development of targeted assays. More recently, parallel reaction monitoring (PRM) has become a promising alternative to SRM. In PRM, the entire product ion spectrum is measured, which enables the confirmation of the identity of an analyte, and the refinement of product ion transitions post-acquisition to remove those with interference. Additionally, PRM only requires the selection of a precursor *m/z* and charge for monitoring, simplifying the development of these assays compared to SRM. Because the selection of specific transitions to target is not needed in PRM, the development of a targeted assay is simplified compared to SRM methods.

Despite the promise of targeted proteomics, there have been limitations in the number of peptides that can be measured due to constraints on the number of concurrent analytes that can be measured at any point in time. As the number of concurrent precursors increases, the time spent on each target decreases, sacrificing the sensitivity and precision of the measurement. In an effort to increase the number of targets that can be measured, data independent acquisition (DIA) has emerged as an alternative strategy where relatively wide MS/MS isolation windows are acquired spanning a predefined mass range. This acquisition results in a decrease in the number of scan events necessary per cycle. However, this intentional multiplexing of the MS/MS acquisition results in spectra that are highly chimeric and comes at the expense of selectivity and dynamic range.

Another way to increase the number of peptide targets is to incorporate retention time scheduling into LC-MS/MS acquisition method. It is common to monitor each target for 2 to 5 min, despite peptide peak widths of <20 s. This inefficient scheduling is necessary because of chromatographic shifts that occur over the lifespan of an HPLC column. Irreproducible retention time is particularly pronounced in the nanoliter to microliter per minute flow rates. Recently we reported a prototype data acquisition strategy where the instrument automatically assessed the retention time relative to a reference analysis and adjusted the instrument target list to only sample a peptide in the time immediately prior and after the peptide was expected^8^. This was accomplished while accommodating non-linear and irregular shifts in retention time.

While the power of targeted assays is undeniable, the development of robust methodology for the selection of peptides to use as a proxy of a translated gene product is not straightforward. Due to differences in their inherent physicochemical properties, equimolar peptides of different amino acid sequences can have drastically different responses in a mass spectrometer. This development often relies on the use of stochastically sampled protein digests using data dependent acquisition (DDA) to determine which peptides should be targeted. This strategy is predicated on the assumption that the peptides most frequently identified in DDA experiments will produce peptides with the optimal signal-to-noise ratios for a targeted proteomic experiment – unfortunately, this is not the case^9,10^. There are numerous reasons why a peptide may not be selected for MS/MS during a DDA experiment. Therefore, a peptide that is not observed in this type of experiment should not be excluded in a targeted experiment. Conversely, a peptide that is routinely sampled in a DDA style experiment might not necessarily be a suitable peptide in a targeted analysis. Further challenges exist, such as the determination of when they elute from the LC in a given sample matrix and selected chromatographic setup, and which precursor-to-product transitions should be used for SRM acquisition and analysis, or for PRM data analysis.

Our lab has shown that the use of gas phase fractionation (GPF) with DIA-MS with narrow precursor isolation windows is a powerful strategy for assessing which peptides can be measured well in a DIA experiment with less specific precursor isolation. More recently, we have shown that these DIA based chromatogram libraries are powerful for providing information about 1) which peptides are detectable, 2) the relative abundance of the fragment ions, 3) the retention time and chromatographic characteristics of each peptide, and 4) peptide stability – significantly expediting the targeted assay development process. This original strategy was used to streamline the development of SRM assays but this process is also relevant for PRM^11^.

Here we describe the application of the Thermo Fisher Scientific Stellar mass spectrometer for the implementation of highly multiplex targeted assays in both human CSF and plasma. The Stellar instrument hardware is a combination of a quadrupole mass filter, an ion routing multiple, and a dual pressure, radial ejection, linear ion trap (LIT). This hardware is conceptually similar to the Tribrid series instruments without the Orbitrap analyzer. Additionally, as with the Tribrid instruments, the Stellar can pipeline the quadrupole isolation and analyte fragmentation in parallel with the spectrum acquisition in the LIT. A unique capability of the Stellar is the implementation of an intelligent data acquisition strategy that can accommodate chromatographic retention time shifts by updating the scheduled target list using a real-time retention time alignment as described previously^8^ and now called Adaptive RT. Furthermore, the workflow for developing targeted PRM assays using GFP DIA data has been improved with a Skyline external tool called PRM Conductor.

## RESULTS

### Scheduling LIT PRM assays with adaptive real-time alignment

The Stellar mass spectrometer was configured to collect PRM data using its Adaptive RT algorithm. The Adaptive RT algorithm has been described in detail previously^8^. In brief, reference spectra from DIA spectra with 50 *m/z* wide isolation windows can be compressed and serialized to a file having the extension .rtbin. This file can be selected and embedded into targeted MSn instrument methods, which causes the method editor to include those same DIA acquisitions into the assay. During execution of the targeted MSn assay, the data from the DIA acquisitions are compared with the embedded data in real-time to estimate retention time shifts compared to the reference and to update the set of active targets at a period of 3.5 times per LC peak width. Creation of the .rtbin file and embedding into a method can be automated with the PRM Conductor program.

### A generalized workflow for the development of reproducible, highly multiplex protein PRM assays

Previous work from our group has demonstrated the use of gas-phase fractionation (GPF) data-independent acquisition (DIA) for the assessment of the peptides and proteins detectable in a given sample matrix^12–14^. The information acquired in these libraries can then be used to inform the selection of precursors for targeted assays^11^. We use GPF-DIA libraries both acquired with the Orbitrap (OT) analyzer of an Exploris 480 mass spectrometer or with the linear ion trap (LIT) of a Stellar mass spectrometer to show the feasibility of generating assays that are selected from and aligned to DIA data from either an OT system or from the same Stellar MS LIT used for the PRM acquisition. We generated two types of assays based on the GPF DIA data. A neurodegenerative disease protein assay was constructed to measure all detectable peptides derived from a set of proteins informed by both literature and previous experimental results in CSF. Another PRM assay was constructed for measuring the maximum reproducibly sampled precursors in Mag-Net enriched plasma EVs in dementia.

### Generation and scheduling of PRM assays using gas-phase fractionated libraries

The GPF library acquired with 4 *m/z* overlapping windows from the OT did result in a larger number of peptides and proteins detected when compared to the LIT library searched with MSFragger. For the precursors that were detected in both systems there is a strong correlation in the observed retention times (Figure 2a).

**Figure 1.**
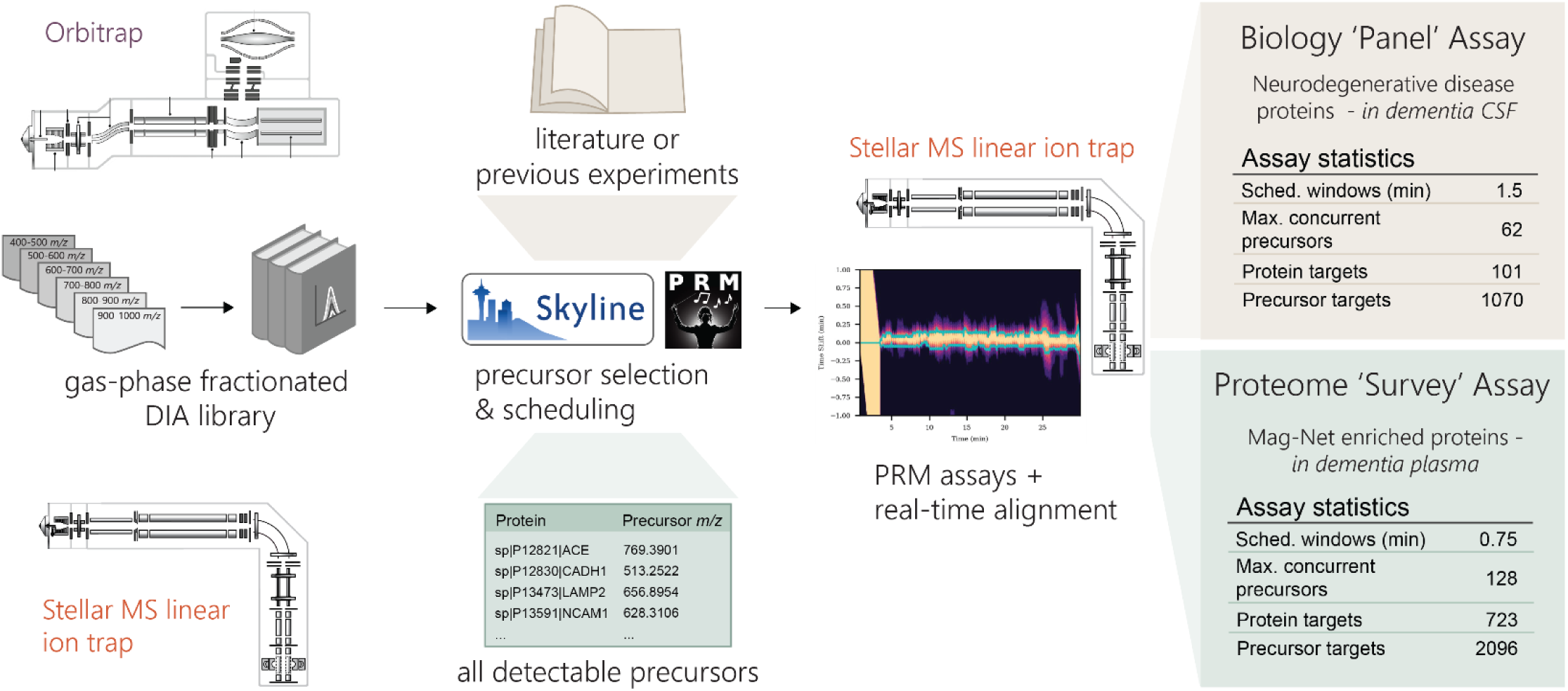
Workflow for the generation of scheduled PRM assays with gas-phase fractionated (GPF) DIA libraries. GPF-DIA libraries can be acquired either directly on a Stellar MS linear ion trap or by an Orbitrap instrument, and their results extracted in Skyline. With the use of the PRM conductor tool in Skyline precursors can be refined for chromatographic performance and signal, and precursors can be selected to optimize both the selection of precursor *m/z* and relative retention time. Selected precursors are then acquired by PRM on the Stellar MS with dynamic real-time alignment. We demonstrate the selection of precursors for an assay targeting a panel of neurodegenerative associated proteins in CSF, and the selection of precursors for a ‘survey’ assay of the measurable proteome of Mag-Net enriched plasma.

**Figure 2.**
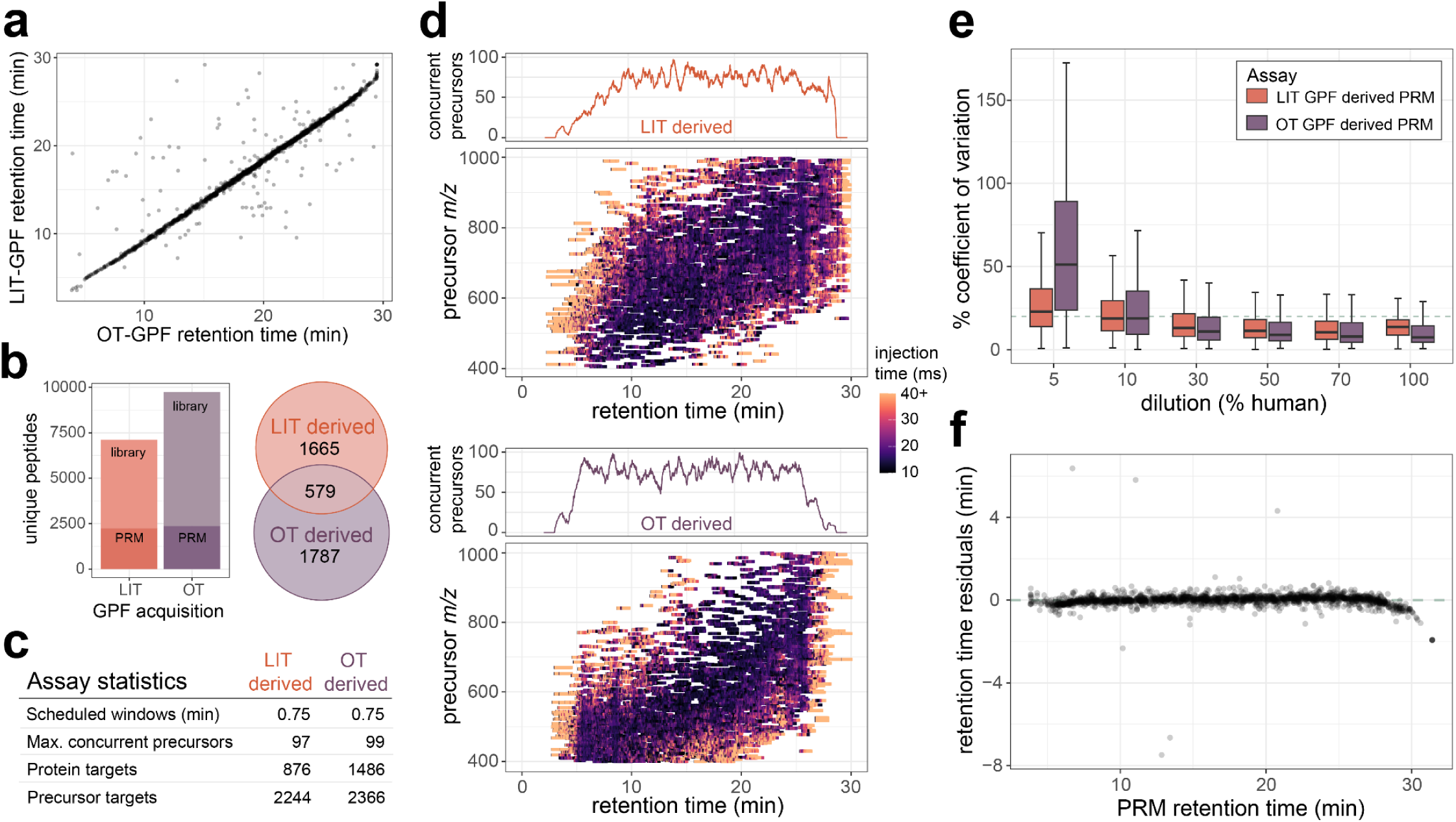
CSF survey PRM assays developed from either Stellar MS linear ion trap (LIT) GPF library or Orbitrap™ (OT) GPF library data. a) Gas-phase fractionated DIA was acquired by Exploris Orbitrap (OT) with 4 *m/z* staggered windows, or by Stellar MS linear ion trap (LIT) with 2 *m/z* discrete windows. From both libraries precursor targets were selected for LIT PRM assays using the PRM Conductor tool to filter and maximize selection of the number of precursors. b) General characteristic of the two assays generated from either GPF DIA library, with the number of concurrent precursors across the retention time for each assay. c) Targeted precursor acquisition windows from PRM conductor to optimize the windows across the precursor mass range and retention time space for both assays. e) The % coefficient of variation from peptides targeted in both assays at 5%, 10%, 50%, and 100% human CSF dilutions in chicken serum. f) Residuals from predicted retention times based on the OT GPF library and the measured retention times on the Stellar PRM assay derived from the OT GPF library.

From these two libraries two separate assays were constructed to measure as many precursors or proteins as possible while maintaining an average of 6 points across the peak for 13.9 second peak average. For the assay generated from the LIT GPF a total of 2,248 precursors mapping to 876 proteins were targeted, while the assay generated from the OT GPF targeted a total 2,367 precursors mapping to 1,486 proteins. Both assays targeted a different portion of the proteome, with the assay generated from the LIT GPF library prioritizing the number of precursors and the assay generated from the OT GPF library prioritizing the number of proteins, while also maintaining a similar number of precursors (Figure 2b-c).

The precursors selected from the LIT GPF are distributed evenly both across chromatographic separation and precursor *m/z*. The precursors selected from the OT GPF were also distributed evenly across retention time, while enriched for precursors in the lower precursor *m/z* range. For precursors targeted in both PRM assays the OT GPF based assay has slightly lower median %CV for the least dilute points on a matrix-matched calibration curve (Figure 2d). At around 10% human CSF in the chicken serum background, both assays have similar medians, while lower concentrations have worse reproducibility in the OT GPF based assay (Figure 2e). PRM acquired on the Stellar MS system has a strong correlation in retention times to the OT GPF library detections with the same separation conditions (Figure 2f). An initial assay consisting of 4 separate injections with 2 min windows were compared to the original OT GPF data, which ensured accurate alignment and scheduling for the 0.75 minute scheduled single injection method.

### Highly multiplex LIT PRM has similar quantitative capacity in CSF as orbitrap DIA

To benchmark the quantitative performance of highly multiplex assays on the Stellar MS we compared a matrix-matched calibration curve of cerebrospinal fluid (CSF) into chicken serum acquired by 12 *m/z* staggered window DIA (6 *m/z* precursor specificity after demultiplexing) on the Exploris 480 OT and by PRM on the Stellar MS. The PRM assay was constructed for 2,244 precursors detected in a LIT GPF library as described above. For peptides measured in both experiments there was a similar distribution in the calculated limits of quantitation, with the DIA median 8.5% and PRM median 9.7% (Figure 3a). The reproducibility of the PRM was better at lower dilutions on the calibration curve compared to the same peptides measured by DIA. The accuracy of the quantification was assessed by calculating the log2 ratio of the diluted sample peptide abundance by the undiluted sample peptide abundance. The observed ratio distributions for the PRM assay are tightly centered around the expected ratios for the higher % human samples, with the distribution widening at lower concentrations (Figure 3c).

**Figure 3.**
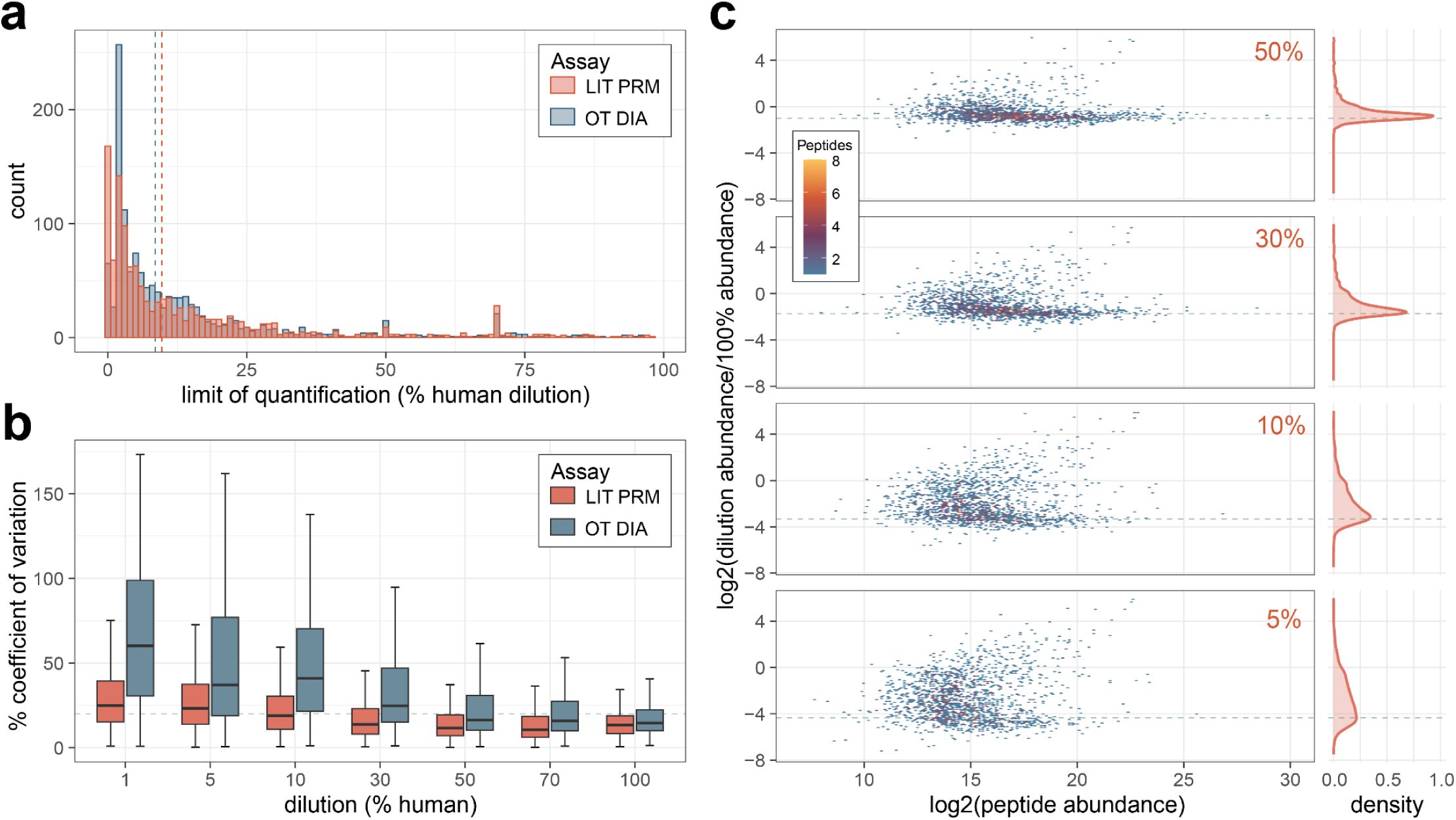
Comparison of quantification by Stellar MS PRM and Orbitrap (OT) DIA. Calibration curves of cerebrospinal fluid diluted into chicken serum were used to construct figures of merit. All data shown are for peptides measured on both Stellar MS LIT PRM assay and Exploris OT DIA experiment with 6 *m/z* precursor windows after demultiplexing. a) The distribution of the limit of quantification after transition optimization for both systems. b) Coefficient of variation from triplicate curve measurements. c) Distribution of peptide ratios at points on the calibration curve acquired by Stellar MS LIT PRM, ratios are calculated as the log2 ratio of mean summed precursor area of the dilution to the 100%. Dashed lines indicate the expected ratios for each dilution point.

### PRM assay size and quantification in Mag-Net enriched plasma EV particles

While the number of precursors targeted is a qualitative metric of the assay, we wanted to examine the quantitative accuracy of these large assays. Two assays were constructed based on a GPF-DIA library with 1

*m/z* windows on Mag-Net enriched human plasma EVs. For the large assay 3,501 precursors mapping to 1,027 proteins were targeted with 0.75 minute scheduled windows, resulting in a maximum of 174 concurrent precursors on a 30 minute gradient. For the small assay 1,599 precursors mapping to 735 proteins were targeted with 0.75 minute scheduled windows, resulting in a maximum of 98 concurrent precursors on a 30 minute gradient (Figure 4a). Both assays were used to measure in triplicate a matrix-matched calibration curve constructed from digested Mag-Net enriched EVs prepared from human plasma diluted into chicken plasma.

**Figure 4.**
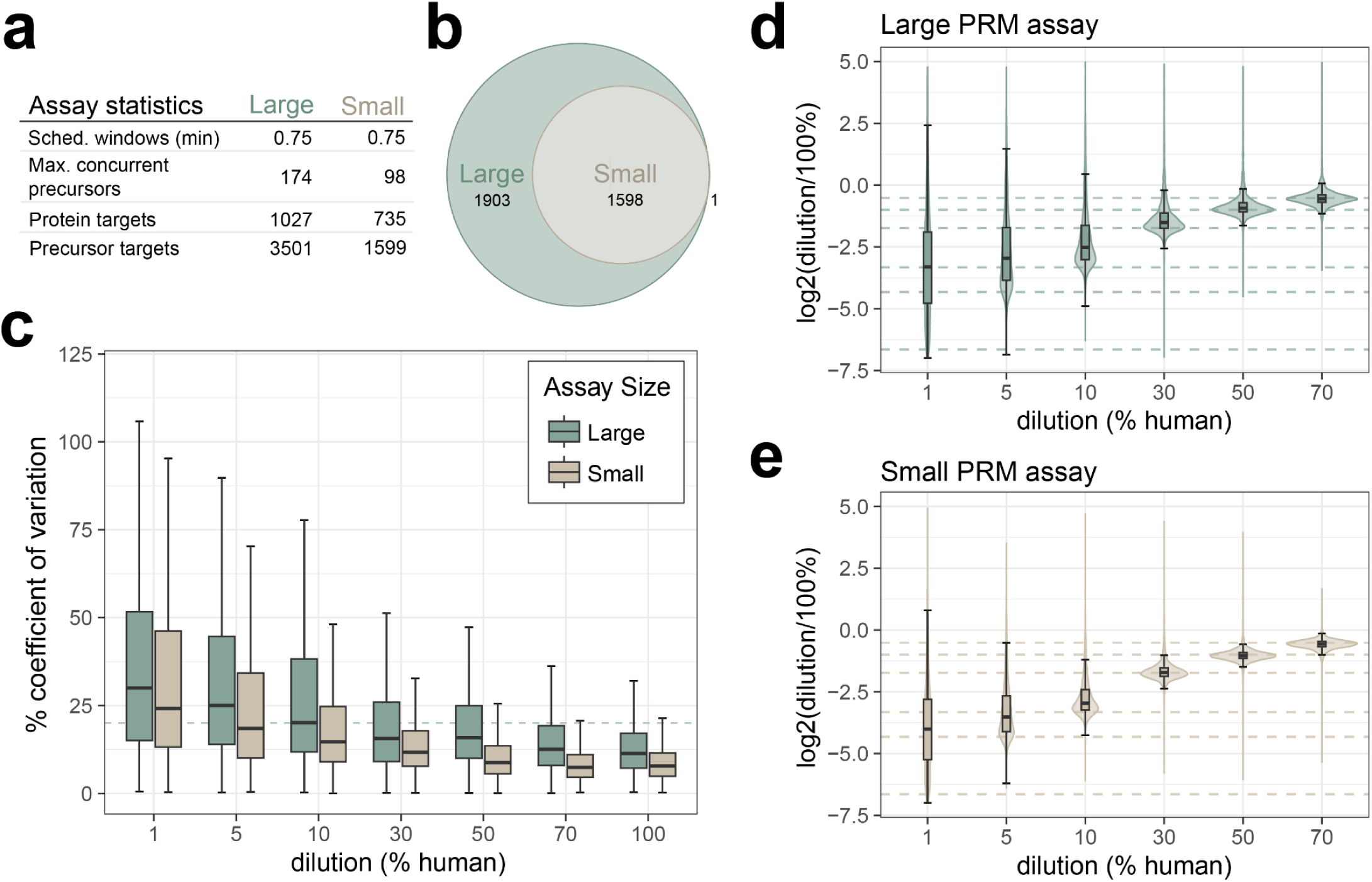
Quantitative performance of Mag-Net enriched plasma EV PRM survey assay. a) A GPF DIA library was acquired on the Stellar mass spectrometer, and two separate assays were constructed for either 3501 or 1599 precursors. Assays were acquired on a 30 min gradient, with 0.75 scheduled windows. b) The small assay was a subset of the large assay, with more stringent filtering parameters. Figures of merit were constructed from a matrix-matched calibration curve analyzed by both assays. c) The precision of the assays from 1% to 100% dilutions as measured by % coefficient of variation, with the dashed line indicating a 20% CV. The accuracy of the assay from 1% to 70% dilutions, calculated as the log2 ratio of mean summed precursor area of the dilution to the 100% for the d) large assay and e) small assay. Dashed lines indicate the expected ratios for each dilution point.

**Figure 5.**
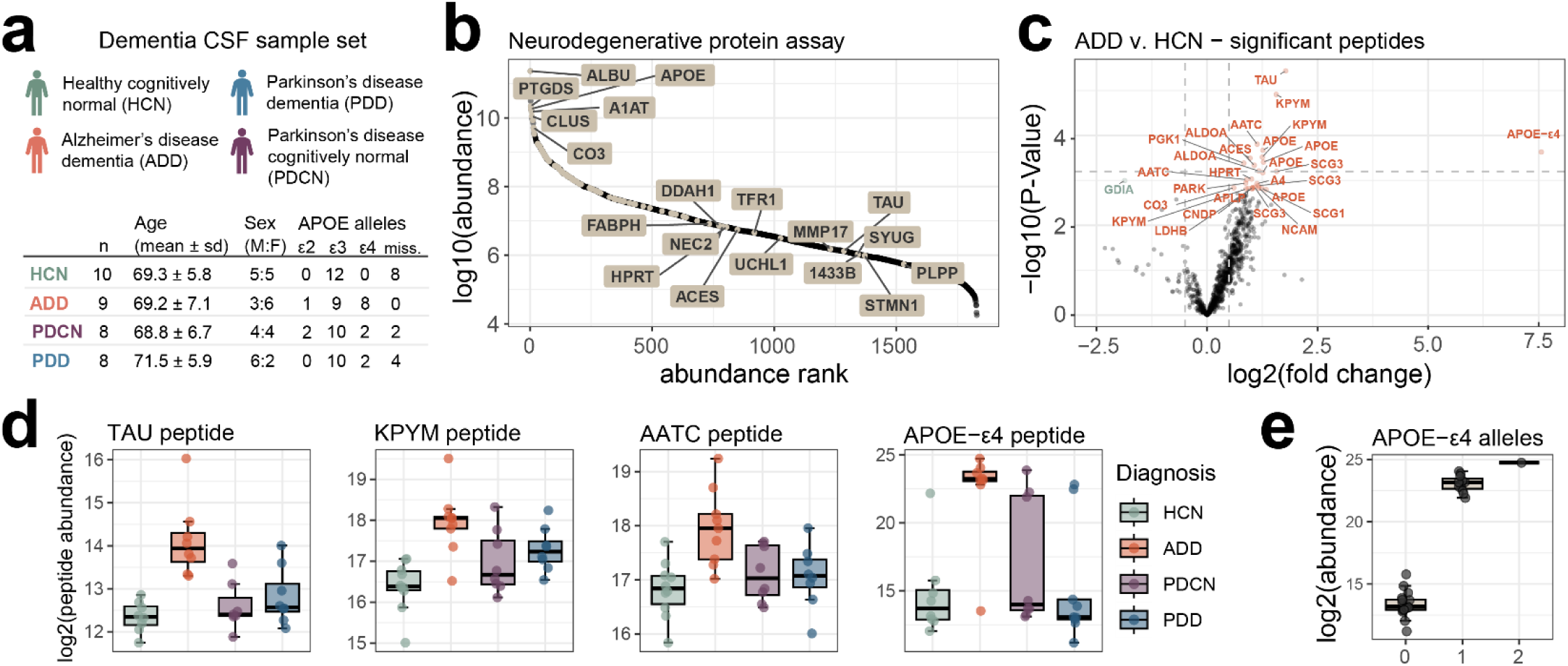
CSF neurodegenerative disease associated protein PRM assay. a) Dementia CSF sample set statistics. b) The distribution of the targeted proteins (highlighted) across the whole CSF dynamic range as determined by orbitrap GPF DIA library. c) Differential abundance of target precursors between samples diagnosed with Alzheimer’s disease (AD) and healthy controls (HC). Precursors are labeled with their corresponding protein ID, and Benjamini-Hochberg adjusted p-value <0.05 colored, with orange representing precursors increased in AD relative to HC, and green decreased in AD relative to HC. d) Select peptides with significantly different abundance in AD compared to the other diagnosis groups. e) APOE-ε4 isoform specific peptide LGADMEDVR abundance separated by the number of APOE-ε4 alleles based on genotyping. The abundance of the peptide with zero alleles is based on the integration of the background signal in the absence of the peptide.

The larger of the two assays was less reproducible across all dilution points, with the median % coefficient of deviation surpassing 20% at the 10% dilution in the large assay (20.2%) and at the 1% dilution in the smaller assay (24.4%) (Figure 4c). The observed ratio distributions are tightly centered around the expected ratios for the higher % human samples for both assays, with the smaller assay distributions still closely centered around the expected ratio down to 5% human dilution (Figure 4d-e).

### Stellar MS PRM assay targeting neurodegenerative disease proteins of interest in CSF

A list of 105 proteins was compiled from previously published targeted assays and discovery proteomics results for Alzheimer’s and Parkinson’s disease characterization^15,16^. From a 60 min, 2 *m/z* window GPF DIA library acquired on the Stellar mass spectrometer we detected peptides mapping to 101 of those proteins. After filtering for signal quality and setting a limit of a 1.7 second cycle time to target approximately 6 spectra per peak we selected 1,070 precursors with window optimization and a minimum of 1.5 minute windows. This resulted in a neurodegenerative disease panel assay with 62 concurrent precursors and dynamic real-time alignment.

The neurodegenerative disease panel assay was used to measure a dementia CSF sample set consisting of individuals diagnosed as both Alzheimer’s disease dementia (AD) and Parkinson’s disease (PD) with and without dementia (PDD and PDCN respectively). Of the peptides measured 29, mapping to 18 proteins, were significantly different between the AD and healthy cognitively normal (HCN) individuals (Figure 4d). The peptide mapping to Tau protein was most significantly changing in AD, which has been previously described (Mann paper). The largest fold change difference was observed in the APOE-ε4 isoform specific peptide LGADMEDVR. APOE-ε4 is found at a higher frequency in individuals diagnosed with AD compared to the general population, making it a risk factor for disease [Nature APOE review paper, newer Altmann paper]. Our sample set recapitulates this observation, with 7 AD individuals having 8 alleles (44% allele frequency), compared to 4 alleles in the rest of the individuals with genotyping data available (19 individuals; 10.5% allele frequency). When the APOE-ε4 peptide abundance is grouped by APOE-ε4 allele status there is a clear trend of abundance matching the genotype. The signal reported for the APOE-ε4 peptide in individuals without the allele is due to the integration of background signal. Because this assay was based on proteins with known association with either Alzheimer’s or Parkinson’s it is reassuring that we find differences despite the relatively small cohort size.

### Mag-Net enriched plasma EV PRM survey assays

For the Mag-Net enriched plasma EV particles, a targeted “survey style” assay was compiled for 2,096 precursors mapping to 723 proteins. Precursors were selected and scheduled based on a GPF 1 *m/z* DIA library collected on a pool of the dementia plasma EV samples. We call the assay “survey style” because it was meant to represent a sampling of peptides that could be measured with high precision and accuracy and not necessarily provide classification between conditions or disease states. The reproducibility of peptide abundances from QC samples included in the experiment was reported previously^17^ using an Orbitrap Eclipse with 12 *m/z* staggered window DIA (6 *m/z* precursor specificity after demultiplexing) – available at https://panoramaweb.org/Mag-Net.url. The PRM survey assay resulted in similar reproducibility for those samples, with highly correlated abundances across QC samples (Figure 6b).

**Figure 6.**
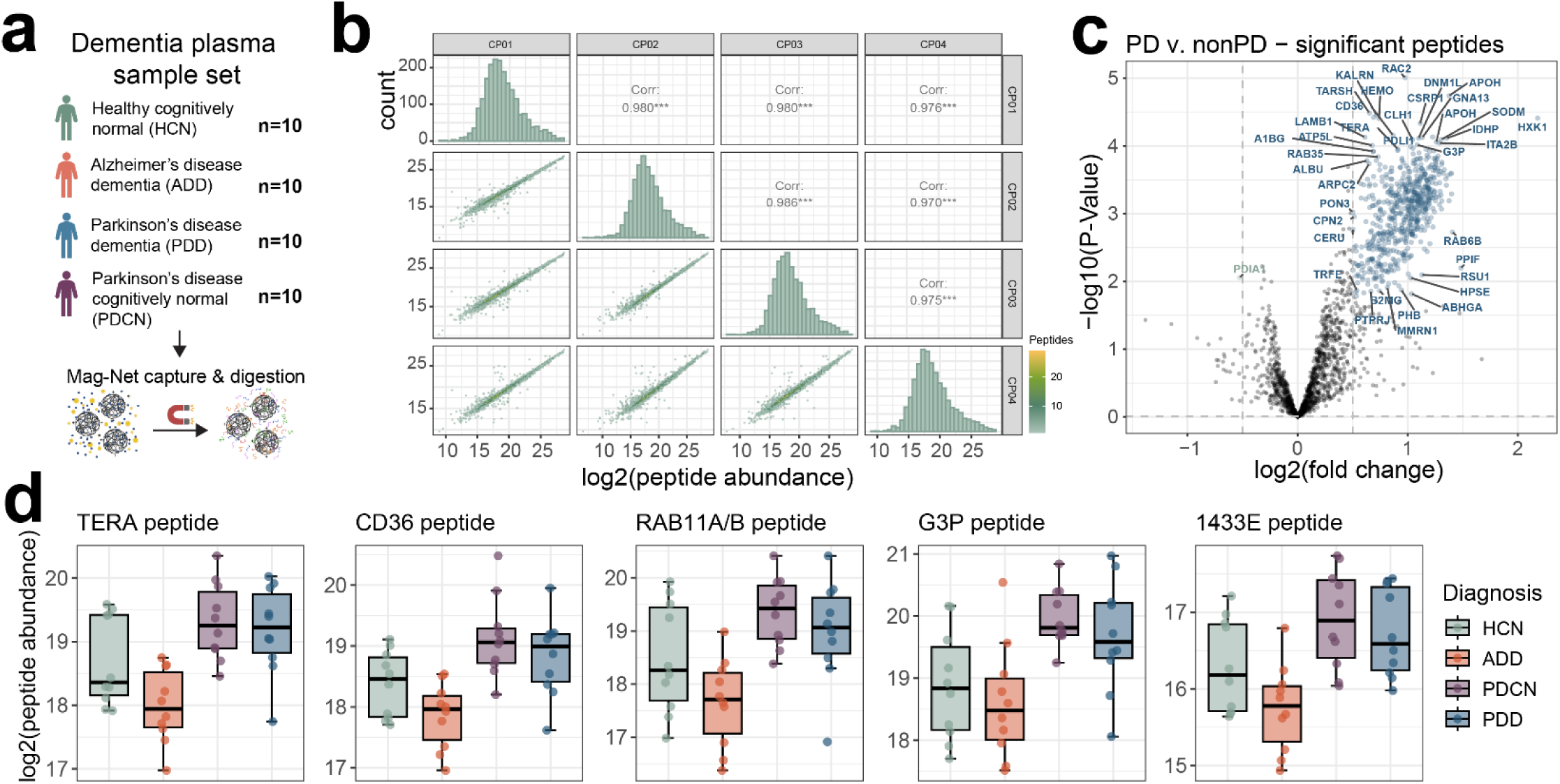
Mag-Net enriched plasma EV survey PRM assay captures distinct differences in Parkinson’s disease. a) A plasma sample set consisting of age-matched healthy cognitively normal, Alzheimer’s disease dementia, Parkinson’s disease dementia, or Parkinson’s disease cognitively normal was enriched for EVs with the Mag-Net method and analyzed by PRM survey assay. b) Peptide abundance correlations and distribution between sample preparation QC replicates. c) TIC normalized peptide abundances between Parkinson’s with and without dementia compared to both healthy cognitively normal and Alzheimer’s disease dementia, with significantly changing peptides highlighted in blue if increased in Parkinson’s and green if increased in non-Parkinson’s samples. Peptides are labeled with their corresponding protein identifier. d) A selection of significantly changing peptides from proteins previously found to be differential in Parkinson’s disease studies.

Between individuals diagnosed as Parkinson’s and those that are not Parkinson’s, a total of 671 peptides mapping to 315 protein groups have significantly altered abundance (Figure 6c). This includes peptides mapping to the proteins TERA (also known as VCP), CD36, RAB11A/RAB11B, G3P (aka GAPDH), and 1433E (aka YWHAE) which have previously been associated with Parkinson’s disease^18–25^ (Figure 6d). Based on gene ontology enrichment analysis of the proteins with significantly changing peptides, there is an enrichment for proteins involved in Platelet Aggregation (GO:0070527) and Homotypic Cell-Cell Adhesion (GO:0034109).

## DISCUSSION

Neurodegenerative diseases, such as Alzheimer’s and Parkinson’s disease, often begin years before noticeable symptoms appear. We are currently in dire need for robust and reliable assays that can monitor disease progression and also assess treatment effectiveness. Biofluids play an important role in being able to quantify proteins that are predictive of disease specific neurodegeneration. Two of the most important biofluids are blood plasma and cerebrospinal fluid as they provide a relatively non-invasive way to sample proteins for use as biomarkers that can aid in the detection, prognosis, and monitoring of the progression of disease. Here we demonstrate the development of highly multiplex targeted assays in both CSF and from enriched plasma EVs that were implemented in a few days of instrument time using a cost-effective quadrupole-radial ejection linear ion trap mass spectrometer – the Stellar.

We demonstrate the value of using DIA data collected using GPF to create a library that can inform the development of highly multiplex targeted peptide PRM assays in CSF and plasma. GPF is powerful because the method has a similar selectivity (e.g. similar quadrupole precursor isolation width) and sensitivity (based on the injection time) as a fully targeted data acquisition. GPF normally comes at the expense of requiring multiple injections of the same sample to cover the *m/z* range of interest with sufficiently narrow DIA windows and long enough injection times. This enables assays to be developed directly within the sample matrix and the resulting library can be mined to create assays for different purposes^11^. Furthermore, we make use of a new Skyline external tool, PRM Conductor, that can select peptides representative of proteins not only based on traditional mass spectrometry and chromatographic performance metrics but also based on the optimal use of the entire chromatographic separation.

We show that it is feasible to use libraries generated using GPF on either high resolution accurate mass (HRAM) Orbitrap platforms or directly using the nominal mass resolution Stellar MS. The Adaptive RT alignment performs well between instrument platforms and with data generated in different laboratories. That said, while the library generated using the Orbitrap platform was significantly larger than the library generated using Stellar MS, the assay generated using nominal mass GPF DIA data performed better quantitatively on the Stellar MS. This suggests that using a HRAM instrument to inform a targeted assay with a nominal mass resolution linear ion trap might not always be ideal. It is likely that peptides selected from Orbitrap data might not be as selective or have a similar dynamic range on the nominal mass resolution Stellar MS. Thus, selecting peptides using the GPF data on the Stellar MS will likely have greater success in a targeted PRM experiment using a Stellar MS.

The generation of highly multiplex assays enable us to measure more analytes of interest in a targeted manner. We demonstrate the use of an assay for the measurement of 101 proteins of interest in CSF, targeting 100% of the confidently detected peptides from a GPF library. While many peptides from the same protein change in the same direction between conditions, our ability to target many peptides per protein increases the chance of detecting individual peptides that change significantly between groups. We believe these large assays will be important in the validation or confirmation of many biologically interesting proteins in a single assay with minimal effort. With the large PRM assays in the Mag-Net enriched plasma EVs, we present a use case for a targeted assay that is not informed by a specific biological hypothesis. By selecting for reproducibly measured precursors while maximizing the number of precursors we target across a gradient and across the precursor range, we are still able to capture interesting biology in a dementia plasma cohort. We believe these highly multiplex quantitative assays that monitor biomedically relevant proteins will provide a much needed bridge from discovery experiments to large scale clinical studies.

## METHODS

### Cerebrospinal fluid and plasma samples

Both lumbar cerebrospinal fluid (CSF) and plasma in EDTA anticoagulant were collected for age and sex matched samples from four diagnosis categories: Parkinson’s disease cognitively normal (PDCN), Parkinson’s disease with dementia (PDD), Alzheimer’s disease dementia, and healthy cognitively normal (HCN). Diagnostic criteria for ADD and PD and the assessments for the cognitive status determination have previously been described^26^. Study protocols were approved by the institutional review board of Stanford University and written informed consent was obtained from all participants or their legally authorized representative.

### Cerebrospinal fluid protein digestion

A 25 µg aliquot of each human CSF sample was diluted to 1% SDS, 50 mM Tris pH 8.5, containing 400 ng yeast enolase protein added as a process control. The sample was reduced with 10 mM TCEP and the free thiols alkylated with 15 mM iodoacetamide in the dark for 30 min. The remaining iodoacetamide was quenched with the addition of 10 mM DTT for 15 min. Proteins were prepared using protein aggregation capture (PAC) and trypsin digestion as described previously^27,28^. Briefly, 130 µg of MagResyn Hydroxyl beads (ReSyn Biosciences) were added to the reduced and alkylated CSF sample for a 1:5.2 protein:bead ratio. The proteins were aggregated to the beads with the addition of 70% acetonitrile for 10 min. The beads were washed 3 times in 95% acetonitrile and 2 times in 70% ethanol for 2.5 minutes each using a ThermoFisher Scientific KingFisher Flex. Peptides were released from the beads by adding the beads to 100 mM ammonium bicarbonate with trypsin (enzyme substrate ratio 1:20). The digestion was quenched by acidification to 0.5% trifluoroacetic acid and spiked with Pierce Retention Time Calibrant (PRTC) peptides to a final concentration of 50 fmol/µl.

### Plasma extracellular vesicle enrichment and digest using Mag-Net

Extracellular vesicles (EVs) were enriched from plasma samples and digested using a ThermoFisher Scientific KingFisher Flex as described previously^17^. Briefly, 100 µl of plasma was combined 1:1 with Binding Buffer (100 mM Bis-Tris Propane, pH 6.3, 150 mM NaCl). MagResyn strong anion exchange (SAX, ReSyn Biosciences) were first equilibrated 2 times with 500 µl Equilibration/Wash Buffer (50 mM Bis-Tris Propane, pH 6.5, 150 mM NaCl) with gentle agitation and then combined in a 1:4 ratio (volume beads:volume starting plasma) with the plasma:binding buffer sample for 45 min at room temperature. The beads were washed with 500 µl Equilibration/Wash Buffer 3 times for 5 min with gentle agitation. The enriched membrane particles on the beads were then solubilized and reduced in 100 µl 50 mM Tris, pH 8.5/1% SDS/10 mM Tris (2-carboxyethyl) phosphine (TCEP) at 37°C and gentle agitation with 800 ng enolase standard added as an internal quality control [QC Preprint]. Following reduction, the plate was removed from the Kingfisher Flex.

Samples were alkylated with 15 mM iodoacetamide in the dark for 30 min and then quenched with 10 mM DTT for 15 min. The samples were processed using PAC with minor modifications^27,28^. Briefly, the samples were adjusted to 70% acetonitrile, mixed, and then incubated for 10 min at room temperature to aggregate proteins onto the SAX bead surface. The SAX beads were washed 3 times in 95% acetonitrile and 2 times in 70% ethanol for 2.5 min each. Samples were digested for 1 h at 47°C in 50 mM Tris, pH 8.5 with Thermo Scientific Pierce porcine trypsin at a ratio of 20:1 protein to trypsin. The digestion was quenched in 0.5% trifluoroacetic acid and spiked with Pierce Retention Time Calibrant peptide cocktail (Thermo Fisher Scientific) to a final concentration of 50 fmol/μL. Peptide digests were stored at −80°C.

### Matrix-matched calibration curve preparation

Matrix-matched calibration curves were made for both CSF and Mag-Net enriched plasma EV samples^29^. For the CSF curve, a 10% diluted chicken serum sample was used as the matched matrix. For the Mag-Net enriched plasma EV samples, a pool of human plasma was diluted into chicken plasma (Innovative Research, Novi, MI). For both the CSF and Mag-Net enriched plasma EV samples, a calibration curve was made using volumetric human:chicken percentages or 100%, 70%, 50%, 30%, 10%, 7%, 5%, 3%, 1%, 0.7%, 0.5%, 0.3%, and 0.1%.

### CSF survey PRM assays and corresponding DIA

Gas-phase fractionated (GPF) DIA libraries were acquired both on an Exploris 480 Orbitrap and Stellar mass spectrometer linear ion trap. CSF peptide digest was separated with a 30 minute gradient from Mobile phase A of 5% (0.1% Formic Acid in water) to 55% Mobile phase B (1% Formic acid in 80% Acetonitrile) on a 0.5 cm, 300 µm inner diameter trap column with a 15 cm 150 um inner diameter C18 PepSep separations column with a Vanquish Neo UHPLC. Orbitrap GPF DIA data was acquired with an Exploris 480 instrument using 4 *m/z* staggered windows from precursor mass range 400-1000 *m/z*. Separate injections were performed to measure each 100 *m/z* precursor mass range (400-500 *m/z*, 500-600 *m/z*, 600-700 *m/z*, 700-800 *m/z*, 800-900 *m/z*, 900-1000 *m/z*), with NCE of 27, 1000% normalized AGC target, and an orbitrap resolution of 30,000. Wider isolation window DIA was performed with an Exploris 480 for the 400-1000 *m/z* precursor mass range, with 12 *m/z* staggered windows with NCE of 27, 1000% normalized AGC target, and an orbitrap resolution of 30,000. Raw files were demultiplexed and converted to mzML with MSConvert v3.0.20070, and searched with EncyclopeDIA v2.12.30 with a Prosit library generated from a uniprot canonical human fasta (09/2022) with APOE isoforms and yeast enolase protein sequence added. Exploris 12 *m/z* DIA matrix-matched data were subsequently with EncyclopeDIA and the GPF chromatogram library^14^. All searches were performed using a uniprot canonical human fasta (09/2022) with APOE-ε2 and APOE-ε4 isoform peptides and yeast enolase protein sequence (P00924) added as the background proteome, with normal target/decoy, trypsin enzyme, CID/HCD fragmentation, 10 ppm precursor, fragment, and library mass tolerance, using Percolator v3.01, and a minimum of 3 transitions with no interference. Data were extracted in Skyline-Daily v23.1.1.503 where protein parsimony was performed, and non-unique peptides removed.

Linear ion trap GPF DIA data was acquired with a Stellar mass spectrometer using 2 *m/z* discrete windows with optimized placement for the precursor mass range 400-1000 *m/z*, with NCE of 30%, 125 kDa/s scan rate, dynamic AGC target of 1.0e4, and automatic maximum injection time. Stellar MS-DIA data was searched with MSFragger v3.8 and post-processed with Percolator v3.05 within Skyline. The data was searched against the Uniprot canonical human fasta (09/2022) with APOE isoforms and yeast enolase protein sequence added. MSFragger was searched using a precursor tolerance of +/-1 Da and a fragment tolerance of 0.2 Da. Data were extracted in Skyline where protein parsimony was performed, and non-unique peptides removed.

Two assays were generated from the peptide information in either the Exploris 480 Orbitrap acquired GPF library or the Stellar MS linear ion trap GPF library. CSF peptide digest was separated on a 0.5 cm, 300 um inner diameter trap column with a 15 cm 150 µm inner diameter C18 PepSep separations column with a Vanquish Neo UHPLC. PRM methods were constructed with an Adaptive RT experiment, a full MS1 acquisition, and a triggered MS2 acquisition with the target list and tentative retention time window schedule. The retention time alignment MS scan was 350-1250 *m/z* at 200 kDa/s scan rate, 0.01 s per alignment cycle, AGC target of 3.0e4, and an automatic maximum injection time. The MS1 scan was for 200-1500 *m/z*, with an AGC target of 3.0e4. The targeted MS2 were performed with 1 *m/z* isolation windows, NCE of 30%, 125 kDa/s scan rate, RF lens of 30%, dynamic AGC target of 1.0e4, dynamic maximum injection time, 2.15 s cycle time for a estimated 6 points per peak (avg peak width ∼13.9 s), and Adaptive RT dynamic time scheduling using the .rtbin file created by PRM conductor. Precursor targets for each PRM assay were selected with the PRM conductor tool in Skyline.

For the assay generated from the Exploris orbitrap 480 GPF DIA data a set of precursors were selected by filtering for precursors with a minimum absolute area of 1000, a minimum and maximum peak width of 5.2 and 26.1 s, and a minimum of 3 interference-free transitions. To test the transfer of the alignment between the Exploris and Stellar MS setups a preliminary set of scheduling runs were performed on the Stellar MS ion trap with a 125 kDa/s scan rate, MS2 scan range of 200-1500 *m/z*, a target of 6 points across the peak for 13.07 average measured peak widths resulting in 2.18 s cycle times with optimized window widths and a minimum of 2.4 minute windows. Data was extracted and inspected in Skyline, with some manual peak re-integration to match the library dot product and expected relative retention time. A PRM assay with narrow 0.75 min windows were scheduled based on the 2.4 min precursor window PRM data, with a scan rate of 125 kDa/s, minimum target injection time of 15 msec, scan range from 200-1500 *m/z*, expected LC width of 12.9 s, and a minimum of 6 points per peak resulting in a 2.15 s cycle time with optimized window widths and a minimum of 0.75 minute windows. The OT GPF derived PRM assay was reduced from 3587 precursors from 1735 proteins in the preliminary scheduling runs to 2367 precursors from 1486 proteins for the final assay.

For the PRM assay generated from the Stellar MS linear ion trap GPF DIA data a set of precursors were selected by filtering for precursors with a minimum absolute area of 500, a minimum and maximum peak width of 5.2 and 45 s, and a minimum of 3 interference-free transitions. A scheduled PRM assay was generated with a scan rate of 125 kDa/s, minimum target injection time of 15 msec, scan range from 200-1500 *m/z*, expected LC width of 12.93 s, and a minimum of 6 points per peak resulting in a 2.15 s cycle time with optimized window widths and a minimum of 0.75 minute windows.

For the matrix-matched calibration curves an initial acquisition of each PRM assay was performed on a 50% human CSF in chicken serum to help schedule the appropriate retention time alignment with lower concentrations of human CSF in chicken serum. Alignment of the peaks picked in the 50% were manually inspected to ensure accurate peak picking to match the library spectra. After filtering the LIT GPF derived PRM assay was reduced from 2347 precursors from 917 proteins to 2248 precursors from 876 proteins for the final assay. Each CSF dilution (100%, 70%, 50%, 30%, 10%, 5%, 1%, 0.5%, and 0.1%) was acquired in triplicate for both PRM assays.

### CSF neurodegenerative disease protein PRM assay

Peptides were separated on a 15 cm, 75 µID C18 IonOpticks column by a Vanquish Neo UHPLC and analyzed by DIA and PRM with a Stellar mass spectrometer. A GPF library was collected using 2 Th windows for the precursor mass range 300-1100, with each injection spanning 100 *m/z* of the precursor mass range, with NCE of 30%, 125 kDa/s scan rate, dynamic AGC target of 1.0e4, and automatic maximum injection time. GPF DIA was searched with Proteome Discoverer software with Chimerys search algorithm using the inferys_3.0.0_fragmentation prediction model, with parameters for peptides of 7-30 amino acids with charge +2 to +4, up to 2 missed cleavages, 0.7 Da fragment tolerance, dynamic oxidized methionine modification with a maximum of 2 modifications per peptide, searched with a uniprot canonical human fasta (09/2022) with APOE isoforms and yeast enolase protein sequence added. A list of 105 proteins of interest was compiled from previous studies, with 101 having detectable peptides in the 60m LIT GPF DIA library. For these 101 proteins peptides with missed cleavages were removed (except for the PRDX2 protein, where the missed cleavage peptides were the only unique peptides).

PRM conductor was used to generate a scheduled assay for 1070 precursors after filtering for peaks widths with a minimum of 6 seconds and a minimum of 3 transitions with no interference. PRM methods were constructed with an Adaptive RT DIA experiment with twelve 50 Th isolation width acquisitions, followed by a full MS1 acquisition, and a targeted MS2 experiment with the target list. The retention time alignment DIA experiment measured 400-1000 *m/z* with 200 kDa/s scan rate, 0.01 s per alignment cycle, AGC target of 3.0e4, and an automatic maximum injection time. The MS1 scan was for 200-1500 *m/z*, with an AGC target of 3.0e4. The targeted MS2 were performed with 1 *m/z* precursor isolation windows, NCE of 30%, 125 kDa/s scan rate, RF lens of 30%, AGC target of 1.0e4, dynamic maximum injection time, 1.71 s cycle time for a estimated 7 points per peak (avg peak width ∼12 s), and Adaptive RT dynamic time scheduling using the .rtbin file created by PRM conductor. Precursor targets for each PRM assay were selected with the PRM conductor tool in Skyline (1.5 min windows with optimized widths). For the individual dementia CSF samples peaks were manually inspected, and peptides with truncated peaks in >20% of the samples removed from further analysis.

### Mag-Net enriched plasma EV survey PRM assays of different sizes

One µg of digested sample was loaded onto a µPAC HPLC column with a 75 µm internal diameter and 50 cm length. The separation was performed using a flow rate of 350 nl/min using a Vanquish Neo HPLC. Mobile phase A was 0.1% formic acid in water, and mobile phase B was 80% acetonitrile and 0.1% formic acid in water. The gradient ran from 0% B to 30% B in 22.5 min, increased to 45% B at 30 min, increased again to 99% B at 30.25 min, and remained at 99% B until 35.25 min. Stellar Linear ion trap GPF DIA data was acquired with a Stellar mass spectrometer using 1 *m/z* discrete windows with optimized placement for the precursor mass range 400-1000 *m/z*. GPF DIA data were searched with Proteome Discoverer software with SEQUEST and Inferys rescoring.

Two separate PRM assays were constructed: a “large” assay targeting 3501 precursors and a “small” assay targeting 1599 precursors. Precursor targets for the larger assay were selected from the Stellar LIT GPF DIA using PRM conductor, filtering for precursors with a minimum absolute area of 500, a minimum and maximum peak width of 4 and 17 s, and a minimum of 3 interference-free transitions. Precursors in the small PRM assay were selected by filtering targets from the 3501 assay based on their performance in the matrix-matched calibration curve. Specifically, targets were selected for the small assay if they had <20% coefficient of variation and were within 50% of the expected ratio. PRM methods were constructed with an Adaptive RT DIA experiment with twelve 50 Th isolation width acquisitions, followed by a full MS1 acquisition, and a targeted MS2 experiment with the target list. The retention time alignment DIA experiment measured 400-1000 *m/z* with 200 kDa/s scan rate, 0.01 s per alignment cycle, AGC target of 3.0e4, and an automatic maximum injection time. The MS1 scan was for 200-1500 *m/z*, with an AGC target of 3.0e4. The targeted MS2 were performed with 1 *m/z* precursor isolation windows, NCE of 30%, 125 kDa/s scan rate, RF lens of 30%, and dynamic AGC target of 1.0e4, dynamic maximum injection time, and Adaptive RT dynamic time scheduling using the .rtbin file created by PRM conductor.

### Mag-Net enriched plasma EV survey PRM assay for a dementia cohort

One µg of a digested pooled human enriched plasma EV sample was loaded onto a µPAC HPLC column with a 75 µm internal diameter and 50 cm length, and separated with a 30 minute gradient from Mobile phase A of 5% (0.1% Formic Acid in water) to 55% Mobile phase B (1% Formic acid in 80% Acetonitrile) using a Vanquish Neo HPLC. Stellar linear ion trap GPF DIA data was acquired with 1 *m/z* discrete windows with optimized placement for the precursor mass range 400-1000 *m/z.* Stellar GPF DIA data were searched with Proteome Discoverer software with SEQUEST and Inferys rescoring. Existing results from an Eclipse GPF DIA experiment acquired with 4 *m/z* staggered windows and searched with EncyclopeDIA and a Prosit predicted library, were combined with the Stellar GPF DIA results, and a reduced fasta file generated. The Stellar GPF DIA was then searched against the reduced fasta, and results extracted in Skyline.

Precursor targets were selected with PRM Conductor, filtering for a minimum absolute area of 300, a minimum and maximum peak width of 5 and 17 s, and a minimum of 3 co-eluting transitions without interference. A PRM assay was scheduled with 0.8 minute windows and 1.43 s cycle time, resulting in 3 separate injections to cover 4,136 precursors. The assay was acquired in triplicate with the Stellar MS PRM method with an Adaptive RT DIA experiment with twelve 50 Th isolation width acquisitions, followed by a full MS1 acquisition, and a targeted MS2 experiment with the target list. Results from this preliminary assay were used to further filter for precursors with a TIC-normalized % coefficient of variation below 30% across the three replicates. The final assay targeting 2093 precursors with a 1.5 s cycle time was scheduled through PRM Conductor and used to measure all individual dementia plasma EV samples.

### Statistical analysis

PRM and DIA data was extracted in Skyline Daily with their corresponding libraries. For the matrix matched calibration curves the peak integration boundaries were determined with a python script that uses boundaries from the 100% samples to calculate boundaries for the more dilute samples. The limit of detection and limit of quantification is determined with a python script based on previously described methods^29^, with an additional step to select optimal transitions for improving the figures of merit^30^. Additional statistical analysis and generation of figures was performed in R v4.3.1.

### Data availability

Instrument raw files, database search results, and Skyline files are available on Panorama at https://panoramaweb.org/stellar-biofluid-prm.url. Python and R code for statistical analysis and plotting are available on github (https://github.com/uw-maccosslab/manuscript-stellar-biofluid).

## ACKNOWLEDGEMENTS

This work was supported in part by National Institutes of Health grants R24GM141156, U19AG065156, P30AG013280, and U01DK137097.

## REFERENCES

(1) Katz, D. H.; Robbins, J. M.; Deng, S.; Tahir, U. A.; Bick, A. G.; Pampana, A.; Yu, Z.; Ngo, D.; Benson, M. D.; Chen, Z.-Z.; Cruz, D. E.; Shen, D.; Gao, Y.; Bouchard, C.; Sarzynski, M. A.; Correa, A.; Natarajan, P.; Wilson, J. G.; Gerszten, R. E. Proteomic Profiling Platforms Head to Head: Leveraging Genetics and Clinical Traits to Compare Aptamer- and Antibody-Based Methods. Sci. Adv. 2022, 8 (33), eabm5164. 10.1126/sciadv.abm5164.

(2) Ngo, D.; Sinha, S.; Shen, D.; Kuhn, E. W.; Keyes, M. J.; Shi, X.; Benson, M. D.; O’Sullivan, J. F.; Keshishian, H.; Farrell, L. A.; Fifer, M. A.; Vasan, R. S.; Sabatine, M. S.; Larson, M. G.; Carr, S. A.; Wang, T. J.; Gerszten, R. E. Aptamer-Based Proteomic Profiling Reveals Novel Candidate Biomarkers and Pathways in Cardiovascular Disease. Circulation 2016, 134 (4), 270–285. 10.1161/CIRCULATIONAHA.116.021803.

(3) Emilsson, V.; Ilkov, M.; Lamb, J. R.; Finkel, N.; Gudmundsson, E. F.; Pitts, R.; Hoover, H.; Gudmundsdottir, V.; Horman, S. R.; Aspelund, T.; Shu, L.; Trifonov, V.; Sigurdsson, S.; Manolescu, A.; Zhu, J.; Olafsson, Ö.; Jakobsdottir, J.; Lesley, S. A.; To, J.; Zhang, J.; Harris, T. B.; Launer, L. J.; Zhang, B.; Eiriksdottir, G.; Yang, X.; Orth, A. P.; Jennings, L. L.; Gudnason, V. Co-Regulatory Networks of Human Serum Proteins Link Genetics to Disease. Science 2018, 361 (6404), 769–773. 10.1126/science.aaq1327.

(4) Hoofnagle, A. N.; Wener, M. H. The Fundamental Flaws of Immunoassays and Potential Solutions Using Tandem Mass Spectrometry. J. Immunol. Methods 2009, 347 (1), 3–11. 10.1016/j.jim.2009.06.003.

(5) Hoofnagle Andrew N.; Eckfeldt John H.; Lutsey Pamela L. Vitamin D–Binding Protein Concentrations Quantified by Mass Spectrometry. N. Engl. J. Med. 2015, 373 (15), 1480–1482. 10.1056/NEJMc1502602.

(6) Hoofnagle, A. N.; Roth, M. Y. Improving the Measurement of Serum Thyroglobulin With Mass Spectrometry. J. Clin. Endocrinol. Metab. 2013, 98 (4), 1343–1352. 10.1210/jc.2012-4172.

(7) Plubell, D. L.; Käll, L.; Webb-Robertson, B.-J.; Bramer, L. M.; Ives, A.; Kelleher, N. L.; Smith, L. M.; Montine, T. J.; Wu, C. C.; MacCoss, M. J. Putting Humpty Dumpty Back Together Again: What Does Protein Quantification Mean in Bottom-Up Proteomics? J. Proteome Res. 2022, 21 (4), 891–898. 10.1021/acs.jproteome.1c00894.

(8) Remes, P. M.; Yip, P.; MacCoss, M. J. Highly Multiplex Targeted Proteomics Enabled by Real-Time Chromatographic Alignment. Anal. Chem. 2020, 92 (17), 11809–11817. 10.1021/acs.analchem.0c02075.

(9) Stergachis, A. B.; MacLean, B.; Lee, K.; Stamatoyannopoulos, J. A.; MacCoss, M. J. Rapid Empirical Discovery of Optimal Peptides for Targeted Proteomics. Nat. Methods 2011, 8 (12), 1041–1043. 10.1038/nmeth.1770.

(10) Searle, B. C.; Egertson, J. D.; Bollinger, J. G.; Stergachis, A. B.; MacCoss, M. J. Using Data Independent Acquisition to Model High-Responding Peptides for Targeted Proteomics Experiments. Mol. Cell. Proteomics 2015, 14 (7), 1–40. 10.1074/mcp.M115.051300.

(11) Plubell, D. L.; Huang, E.; Spencer, S. E.; Poston, K.; Montine, T. J.; MacCoss, M. J. Data Independent Acquisition to Inform the Development of Targeted Proteomics Assays Using a Triple Quadrupole Mass Spectrometer. bioRxiv May 31, 2024, p 2024.05.29.596554. 10.1101/2024.05.29.596554.

(12) Ting, Y. S.; Egertson, J. D.; Bollinger, J. G.; Searle, B. C.; Payne, S. H.; Noble, W. S.; MacCoss, M. J. PECAN: Library-Free Peptide Detection for Data-Independent Acquisition Tandem Mass Spectrometry Data. Nat. Methods 2017, 14 (9), 903–908. 10.1038/nmeth.4390.

(13) Searle, B. C.; Pino, L. K.; Egertson, J. D.; Ting, Y. S.; Lawrence, R. T.; MacLean, B. X.; Villén, J.; MacCoss, M. J. Chromatogram Libraries Improve Peptide Detection and Quantification by Data Independent Acquisition Mass Spectrometry. Nat. Commun. 2018, 9 (1), 5128. 10.1038/s41467-018-07454-w.

(14) Pino, L. K.; Just, S. C.; MacCoss, M. J.; Searle, B. C. Acquiring and Analyzing Data Independent Acquisition Proteomics Experiments without Spectrum Libraries. Mol. Cell. Proteomics 2020, mcp.P119.001913. 10.1074/mcp.P119.001913.

(15) Watson, C. M.; Dammer, E. B.; Ping, L.; Duong, D. M.; Modeste, E.; Carter, E. K.; Johnson, E. C. B.; Levey, A. I.; Lah, J. J.; Roberts, B. R.; Seyfried, N. T. Quantitative Mass Spectrometry Analysis of Cerebrospinal Fluid Protein Biomarkers in Alzheimer’s Disease. Sci. Data 2023, 10 (1), 261. 10.1038/s41597-023-02158-3.

(16) Spellman, D. S.; Wildsmith, K. R.; Honigberg, L. A.; Tuefferd, M.; Baker, D.; Raghavan, N.; Nairn, A. C.; Croteau, P.; Schirm, M.; Allard, R.; Lamontagne, J.; Chelsky, D.; Hoffmann, S.; Potter, W. Z. Development and Evaluation of a Multiplexed Mass Spectrometry Based Assay for Measuring Candidate Peptide Biomarkers in Alzheimer’s Disease Neuroimaging Initiative (ADNI) CSF. PROTEOMICS - Clin. Appl. 2015, 9 (7–8), 715–731. 10.1002/prca.201400178.

(17) Wu, C. C.; Tsantilas, K. A.; Park, J.; Plubell, D.; Sanders, J. A.; Naicker, P.; Govender, I.; Buthelezi, S.; Stoychev, S.; Jordaan, J.; Merrihew, G.; Huang, E.; Parker, E. D.; Riffle, M.; Hoofnagle, A. N.; Noble, W. S.; Poston, K. L.; Montine, T. J.; MacCoss, M. J. Mag-Net: Rapid Enrichment of Membrane-Bound Particles Enables High Coverage Quantitative Analysis of the Plasma Proteome. bioRxiv April 2, 2024, p 2023.06.10.544439. 10.1101/2023.06.10.544439.

(18) Chu, S.; Xie, X.; Payan, C.; Stochaj, U. Valosin Containing Protein (VCP): Initiator, Modifier, and Potential Drug Target for Neurodegenerative Diseases. Mol. Neurodegener. 2023, 18 (1), 1–34. 10.1186/s13024-023-00639-y.

(19) Ioghen, O.; Chițoiu, L.; Gherghiceanu, M.; Ceafalan, L. C.; Hinescu, M. E. CD36 – A Novel Molecular Target in the Neurovascular Unit. Eur. J. Neurosci. 2021, 53 (8), 2500–2510. 10.1111/ejn.15147.

(20) Liu, J.; Zhang, J.-P.; Shi, M.; Quinn, T.; Bradner, J.; Beyer, R.; Chen, S.; Zhang, J. Rab11a and HSP90 Regulate Recycling of Extracellular α-Synuclein. J. Neurosci. 2009, 29 (5), 1480–1485. 10.1523/JNEUROSCI.6202-08.2009.

(21) Avila, C. L.; Chaves, S.; Socias, S. B.; Vera-Pingitore, E.; González-Lizárraga, F.; Vera, C.; Ploper, D.; Chehín, R. Lessons Learned from Protein Aggregation: Toward Technological and Biomedical Applications. Biophys. Rev. 2017, 9 (5), 501–515. 10.1007/s12551-017-0317-z.

(22) Gerszon, J.; Rodacka, A. Oxidatively Modified Glyceraldehyde-3-Phosphate Dehydrogenase in Neurodegenerative Processes and the Role of Low Molecular Weight Compounds in Counteracting Its Aggregation and Nuclear Translocation. Ageing Res. Rev. 2018, 48, 21–31. 10.1016/j.arr.2018.09.003.

(23) Davison, E. J.; Pennington, K.; Hung, C.-C.; Peng, J.; Rafiq, R.; Ostareck-Lederer, A.; Ostareck, D. H.; Ardley, H. C.; Banks, R. E.; Robinson, P. A. Proteomic Analysis of Increased Parkin Expression and Its Interactants Provides Evidence for a Role in Modulation of Mitochondrial Function. PROTEOMICS 2009, 9 (18), 4284–4297. 10.1002/pmic.200900126.

(24) Gispert, S.; Kurz, A.; Brehm, N.; Rau, K.; Walter, M.; Riess, O.; Auburger, G. Complexin-1 and Foxp1 Expression Changes Are Novel Brain Effects of Alpha-Synuclein Pathology. Mol. Neurobiol. 2015, 52 (1), 57–63. 10.1007/s12035-014-8844-0.

(25) Kurz, A.; May, C.; Schmidt, O.; Müller, T.; Stephan, C.; Meyer, H. E.; Gispert, S.; Auburger, G.; Marcus, K. A53T-Alpha-Synuclein-Overexpression in the Mouse Nigrostriatal Pathway Leads to Early Increase of 14-3-3 Epsilon and Late Increase of GFAP. J. Neural Transm. 2012, 119 (3), 297–312. 10.1007/s00702-011-0717-3.

(26) Shahid, M.; Rawls, A.; Ramirez, V.; Ryman, S.; Santini, V. E.; Yang, L.; Sha, S. J.; Hall, J. N.; Montine, T. J.; Lin, A.; Tian, L.; Henderson, V. W.; Cholerton, B.; Yutsis, M.; Poston, K. L. Illusory Responses across the Lewy Body Disease Spectrum. Ann. Neurol. 2023, 93 (4), 702–714. 10.1002/ana.26574.

(27) Hughes, C. S.; Moggridge, S.; Müller, T.; Sorensen, P. H.; Morin, G. B.; Krijgsveld, J. Single-Pot, Solid-Phase-Enhanced Sample Preparation for Proteomics Experiments. Nat. Protoc. 2019, 14 (1), 68–85. 10.1038/s41596-018-0082-x.

(28) Batth, T. S.; Tollenaere, Maxim A. X.; Rüther, P.; Gonzalez-Franquesa, A.; Prabhakar, B. S.; Bekker-Jensen, S.; Deshmukh, A. S.; Olsen, J. V. Protein Aggregation Capture on Microparticles Enables Multipurpose Proteomics Sample Preparation*. Mol. Cell. Proteomics 2019, 18 (5), 1027a–11035. 10.1074/mcp.TIR118.001270.

(29) Pino, L. K.; Searle, B. C.; Yang, H.-Y.; Hoofnagle, A. N.; Noble, W. S.; MacCoss, M. J. Matrix-Matched Calibration Curves for Assessing Analytical Figures of Merit in Quantitative Proteomics. J. Proteome Res. 2020, 19 (3), 1147–1153. 10.1021/acs.jproteome.9b00666.

(30) Heil, L. R.; Remes, P. M.; MacCoss, M. J. Comparison of Unit Resolution Versus High-Resolution Accurate Mass for Parallel Reaction Monitoring. J. Proteome Res. 2021, 20 (9), 4435–4442. 10.1021/acs.jproteome.1c00377.

